# Effects of High-Volume versus High-Load Resistance Training on Skeletal Muscle Growth and Molecular Adaptations

**DOI:** 10.1101/2021.07.01.450728

**Authors:** Christopher G. Vann, Casey L. Sexton, Shelby C. Osburn, Morgan A. Smith, Cody T. Haun, Melissa N. Rumbley, Petey W. Mumford, Brian K. Ferguson, Nathan T. Montgomery, Carlton D. Fox, Bradley A. Ruple, James McKendry, Jonathan Mcleod, Adil Bashir, Ronald J. Beyers, Matthew S. Brook, Kenneth Smith, Philip J Atherton, Darren T. Beck, James R. McDonald, Kaelin C. Young, Stuart M. Phillips, Michael D. Roberts

**Affiliations:** School of Kinesiology, Auburn University, Auburn AL, USA; Duke Molecular Physiology Institute, Duke University School of Medicine, Duke University, Durham, NC, USA; Fitomics, LLC, Pelham, AL, USA; Department of Kinesiology, Lindenwood University, St. Charles MO, USA; Department of Kinesiology, McMaster University, Hamilton ON, CANxs; MRI Research Center, Auburn University, Auburn AL, USA; MRC-ARUK Centre of Excellence for Musculoskeletal Ageing Research, Clinical, Metabolic, and Molecular Physiology, University of Nottingham, Derby, UK; Edward Via College of Osteopathic Medicine - Auburn Campus, Auburn AL, USA

**Author notes:** address correspondence to: Michael D. Roberts, Ph. D. Associate Professor, School of Kinesiology, Director, Molecular and Applied Sciences Laboratory, Affiliate Research Professor, Edward Via College of Osteopathic Medicine – Auburn Campus Auburn University, 301 Wire Road, Office 286, Auburn, AL 36849.

**Keywords:** Higher-load resistance training, higher-volume resistance training, muscle hypertrophy, sarcoplasmic protein, myofibrillar protein

## Abstract

**Aim:** We evaluated the effects of higher-load (HL) versus (lower-load) higher-volume (HV) resistance training on skeletal muscle hypertrophy, strength, and muscle-level molecular markers.

**Methods:** Trained men (n=15, age: 23±3 y; training experience: 7±3 y) performed unilateral lower body training for 6 weeks (3x weekly), where single legs were assigned to HV and HL paradigms. Vastus lateralis (VL) biopsies were obtained prior to study initiation (PRE) as well as 3 days (POST) and 10 days following the last bout (POSTPR). Body composition and strength tests were performed at each testing session, and biochemical assays were performed on muscle tissue after study completion. Two-way within subjects repeated measures ANOVAs were performed on all dependent variables except tracer data, which was compared using dependent samples t-tests.

**Results:** A significant (p<0.05) interaction existed for unilateral leg extension 1RM (HV<HL at POST and POSTPR). Six-week integrated sarcoplasmic protein synthesis (iSarcoPS) rates were higher in the HV versus HL leg, while no difference between legs existed for integrated myofibrillar protein synthesis rates. Main time effects existed for unilateral leg press strength (PRE<POST and POSTPR), knee extensor peak torque (PRE and POST<POSTPR), dual-energy x-ray absorptiometry (DXA)-derived upper leg lean mass (PRE<POST and POSTPR), ultrasound-derived VL thickness (PRE and POSTPR<POST), sarcoplasmic protein concentrations (POST and POSTPR<PRE), and tropomyosin and troponin protein abundances (POST and POSTPR<PRE).

**Conclusions:** With the exception of differences in leg extensor strength and iSarcoPS between legs, our data suggest that short-term (6 weeks) HV and HL training elicit similar hypertrophic, strength, and molecular-level adaptations.

## 1. INTRODUCTION

Skeletal muscle hypertrophy has been defined as an increase in the weight or crosssectional area of muscle^1,2^, with the increased volume of muscle coming from an enlargement of each muscle fiber^3–5^. It is generally recognized that resistance training results in skeletal muscle growth through proportional increases in myofibrillar and sarcoplasmic protein content^6–9^. Myofibril proteins are defined herein as the proteins that make up the rigid structure of muscle (e.g., dystrophin, actinin, titin, nebulin, etc.) as well as contractile proteins (e.g., actin and myosin isoforms). In contrast, sarcoplasmic proteins are involved with signal transduction, energy synthesis, energy breakdown, and other metabolic processes^10^.

Recently, there has been interest regarding whether or not higher-load (HL) versus higher-volume (HV) resistance training elicits differential training adaptations at the macroscopic, molecular, and functional levels. Historically, research has typically suggested that HL training elicits superior increases in strength and muscle fiber hypertrophy compared to lower-load HV training^11^. However, Mitchell and colleagues reported that 10 weeks of HL or HV resistance training led to similar increases in muscle hypertrophy as assessed through MRI and fiber histology^12^. Subsequent literature indicates that both HL and HV training can: i) elicit similar changes in skeletal muscle hypertrophy (assessed through either ultrasound or MRI)^13–17^, and ii) elicit similar strength adaptations^13,18^, although equivocal evidence exists suggesting HL training elicits superior strength adaptations^14,15,17^ Reasons for similar outcomes between HL and HV training could be due to total volume lifted being comparable between paradigms. However, few HV versus HL studies have sought to modulate training loads with the intent of accumulating more training volume during HV conditions.

Our laboratory recently reported that six weeks of extremely HV resistance training decreased the relative abundance of myosin heavy chain and actin protein content per milligram of dry tissue^19^. Our findings, as well as those of others who have reported moderate-to-higher volume resistance training elicits similar molecular adaptations^20–22^, led us to postulate that a disproportionate increase in the sarcoplasmic space relative to myofibril protein accretion may be a training adaptation to HV resistance training^23^. More recently, our laboratory demonstrated that lower volume, higher load resistance training (3-5 sets of 2-6 repetitions at 65-90% 1RM) resulted in a maintenance of type I muscle fiber cross-sectional area (fCSA) while increasing type II fCSA. Additionally, no changes in sarcoplasmic protein concentrations were observed despite a modest but significant decrease in actin protein concentrations^24,25^. While preliminary, these two studies from our laboratory suggest that HV resistance training may facilitate a more robust expansion of non-contractile proteins in myofibers, whereas HL training may promote a proportional increase in myofibril protein accretion with muscle growth. However, no single study to date has examined whether HV versus HL training elicit differential molecular adaptations related to the aforementioned findings.

The purpose of the present study was to elucidate whether six weeks of unilateral HV versus HL lower-body resistance training differentially affected metrics of skeletal muscle hypertrophy, strength, and/or molecular variables assessed from skeletal muscle biopsies sampled from the vastus lateralis (VL). The intent of our study was to ensure HV training achieved more training volume relative to HL training. We hypothesized no differences would exist between HV and HL training when examining changes in VL muscle area assessed via magnetic resonance imaging (MRI), or VL thickness assessed via ultrasound. Additionally, we hypothesized that HL training would elicit superior increases in various indices of strength.

However, we posited HV training would result in increased sarcoplasmic protein concentrations and a concomitant decrease in the relative abundances of contractile proteins, whereas HL training would result in no changes in these markers. Additionally, we hypothesized that the integrated sarcoplasmic protein synthesis (iSarcoPS) rates would be greater in HV versus HL training, whereas integrated myofibrillar protein synthesis (iMyoPS) rates would be greater in HL versus HV training.

## 2. RESULTS

### 2.1 Participant characteristics

Baseline participant characteristics can be found in Table 1. Briefly, 15 college-age males (23±3 years) with an average training age of 7±3 years volunteered for this study. At PRE, participants weighed 89.5±11.6 kg with 69.1±7.4 kg being lean soft tissue mass (LSTM) and 17.3±7.5kg being fat mass, on average. Additionally, participants had an average relative squat to body mass ratio of 1.9× body mass (167±34 kg).

**Table 1.** Participant characteristics

| Variable | Mean ± SD |
| --- | --- |
| Age (years) | 23 ± 3 |
| Height (cm) | 182 ± 8 |
| Weight (kg) | 89.5 ± 11.6 |
| Lean soft tissue mass (kg) | 69.1 ± 7.4 |
| Fat tissue mass (kg) | 17.3 ± 7.5 |
| Fat-free mass index | 20.9 ± 2.2 |
| Est. 1RM Squat (kg) | 167 ± 34 |
| Squat relative to body weight | 1.9 ± 0.4 |

N=15 participants. Abbreviations: Est. 1RM, estimated 1 repetition maximum. All measures taken prior to onset of training intervention.

### 2.2 Training Volume and Strength Metrics

Training volumes and strength metrics are presented in Figure 1. Data for 14 of 15 participants is presented for unilateral leg press and unilateral leg extension one repetition maximums (1RM) due to one participant feeling lower extremity discomfort at POST with these exercises.

**Figure 1.**
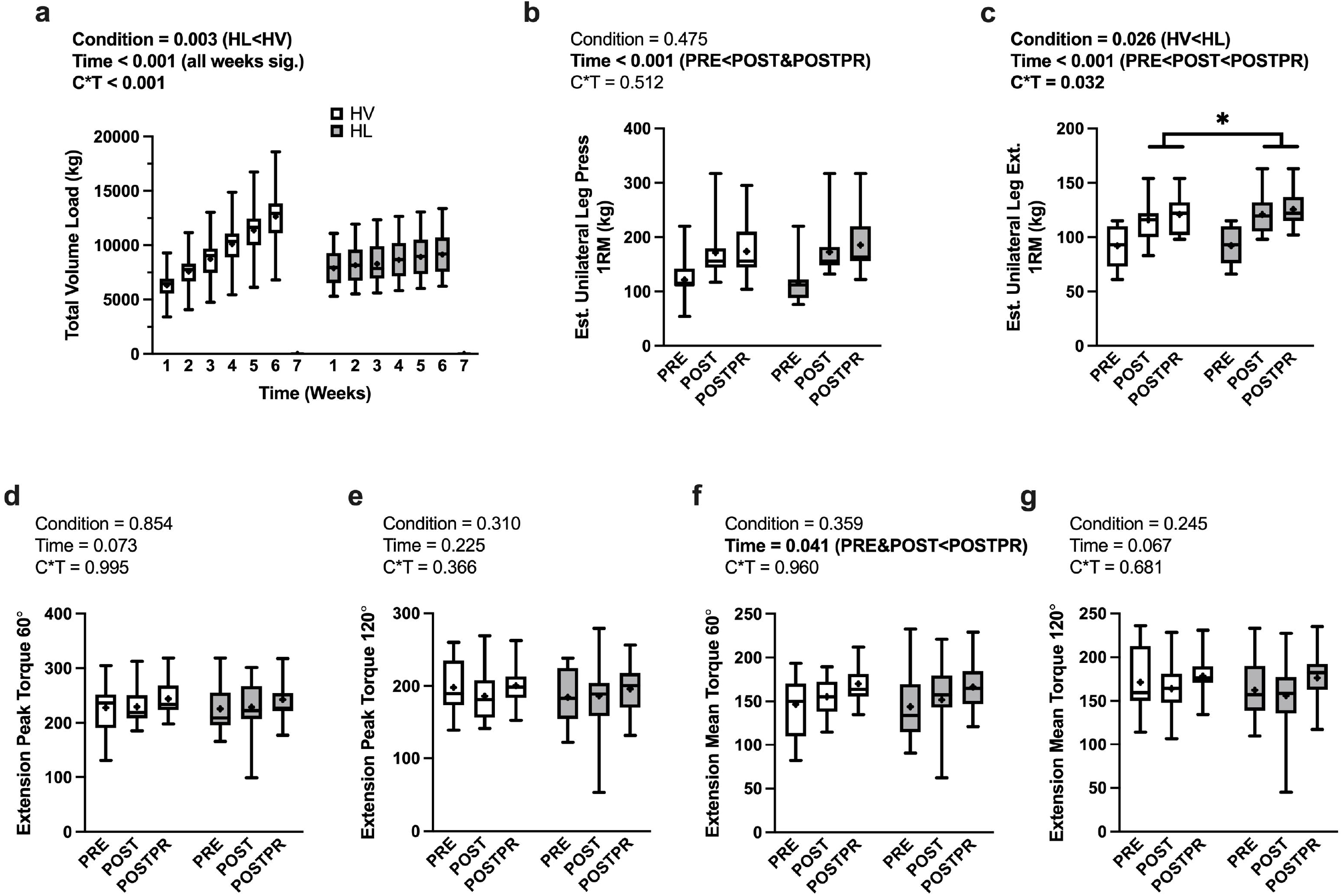
Training Volume and Strength Metrics. Data are presented as box and whiskers plots including median (central horizontal line), 25^th^ and 75^th^ percentile (box), minimum and maximum values (vertical lines) and mean values (cross) for training volume load (panel a), unilateral leg press (panel b), unilateral leg extension (panel c), knee extension peak torque at 60°/s (panel d), knee extension peak torque at 120°/s (panel e), knee extension mean torque at 60°/s (panel f), and knee extension mean torque at 120°/s (panel g). Abbreviations: HV. high-volume; HL, high-load. *, indicates a condition×time interaction whereas POST and POSTPR were greater than PRE.

There was a condition×time interaction observed for lower body training volume (*p*<0.001, η_p_^2^ =0.914; Fig. 1a). Additionally, there was a main effect of condition (*p*=0.003, η_p_^2^=0.467) where the HV condition completed more volume than the HL condition (8100±480 kg versus 7296±421 kg, respectively). Lower body training volume changed over time (*p*<0.001, η_p_^2^=0.955, Fig. 1a) and within each condition over time (HV: *p*<0.001, η_p_^2^=0.952; HL: *p*<0.001, η_p_^2^ = 0.954, Fig. 1a). Post hoc analysis revealed lower training volumes at weeks 1 and 2 in the HV leg compared to the HL leg (*p*<0.001), no differences between conditions at week 3, and higher training volumes in the HV leg at weeks 4-6 as compared to the HL leg (p<0.001).

A condition×time interaction (*p*=0.512, η_p_^2^ =0.050, Fig. 1b) was not observed for estimated unilateral leg press 1RM. Additionally, no main effect of condition (*p*=0.475, η_p_^2^=0.040, Fig. 1b) was observed. There was a main effect of time (*p*<0.001, η_p_^2^=0.818, Fig. 1b) where estimated unilateral leg press 1RM at POST (*p*<0.001) and POSTPR (*p*<0.001) were greater than PRE. A condition×time interaction was observed for estimated unilateral leg extension 1RM (*p*=0.032, η_p_^2^=0.265, Fig. 1c). A main effect of condition (*p*=0.026, η_p_^2^=0.328, Fig. 1c) was also observed where the HL condition (grand mean=113±5 kg) estimated unilateral leg extension 1RM was higher than the HV condition (grand mean=109±5 kg). Estimated unilateral leg extension 1RM also changed over time (*p*<0.001, η_p_^2^ =0.885, Fig. 1c) and within each condition over time (HV: *p*<0.001, η_p_^2^=0.858; HL: *p*<0.001, η_p_^2^=0.884). Post hoc analysis revealed no differences in estimated unilateral leg extension 1RM at PRE; however, the HL condition had higher values at POST and POSTPR compared to the HV condition (p<0.05).

There was no condition×time interaction observed for knee-extensor peak torque at 60°/sec (*p*=0.995, η_p_^2^<0.001, Fig. 1d) and 120° (*p*=0.366, η_p_^2^=0.069, Fig. 1e), or knee-extensor mean torque at 120°/sec (*p*=0.681, η_p_^2^=0.027, Fig. 1g). Additionally, there were no main effects of condition or time observed for the aforementioned variables. Knee-extensor mean torque at 60°/sec changed over time (*p*=0.041, η_p_^2^=0.204, Fig. 1f) where knee-extensor mean torque at 60°/sec was higher at POSTPR than at PRE (*p*=0.029) and POST (*p*=0.043). There were no differences observed between PRE and POST (*p*=0.805).

### 2.3 Body Composition

PRE, POST, and POSTPR whole-body composition changes for all participants are presented in Table 2; notably, these data were derived from dual-energy x-ray absorptiometry (DXA) scans. Total body mass increased over time (*p*<0.001, η_p_^2^ =0.435), where POST (*p*=0.001) and POSTPR (*p*=0.012) body masses were greater than PRE. However, no differences were observed between POST and POSTPR body masses (*p*=0.119). Lean soft tissue mass (LSTM) increased over time (*p*=0.003, η_p_^2^=0.338) where POST LSTM was greater than PRE (*p*=0.002) and POSTPR (*p*=0.014). No significant differences in LSTM were observed between PRE and POSTPR (*p*=0.286). No differences were observed for DXA measured whole body fat mass (*p*=0.097).

**Table 2.** Body composition changes during training

| Variable | PRE | POST | POSTPR | ANOVA |
| --- | --- | --- | --- | --- |
| | Mean ± SD | Mean ± SD | Mean ± SD | p-value |
| Total Body Mass (kg) | 89.3 ± 11.5 | 90.8 ± 11.9 | 90.4 ± 12.2 | **<0.001**<sup>†</sup> |
| DXA Whole Body LSTM (kg) | 69.1 ± 7.4 | 70.2 ± 7.5 | 69.5 ± 7.5 | **0.003** * |
| DXA Whole Body Fat Mass (kg) | 17.3 ± 7.5 | 17.4 ± 7.8 | 17.7 ± 7.8 | 0.097 |

N=15 participants. Abbreviations: DXA, dual energy x-ray absorptiometry; LSTM, lean soft tissue mass. <sup>†</sup>, indicates measurement was higher at POST and POSTPR than PRE ( $p < 0.05$ ); \*, indicates measurement was higher at POST than at PRE and POSTPR ( $p < 0.05$ ).

### 2.4 Segmental Upper Leg Composition

There were no condition×time interactions observed for DXA-derived upper leg mass (*p*= 0.069, η_p_^2^=0.173, Fig. 2a), upper leg LSTM (*p*=0.174, η_p_^2^=0.117, Fig. 2b), or upper leg fat mass (*p*=0.959, η_p_^2^ =0.003, Fig. 2c). A main effect of time was observed for upper leg mass (*p*=0.001, η_p_^2^=0.392, Fig. 2a) where POST (*p*=0.002) and POSTPR (*p*=0.013) were higher than PRE. No differences were observed between POST and POSTPR (*p*=0.240). A main effect of time was observed for DXA upper leg LSTM (*p*=0.001, η_p_^2^=0.418, Fig. 2b) where POST (*p*<0.001) and POSTPR (*p*=0.002) were higher than PRE. No differences were observed between POST and POSTPR (*p*=0.148). No main effects of condition (*p*=0.102) or time (*p*=0.595) were observed for DXA upper leg fat mass.

**Figure 2.**
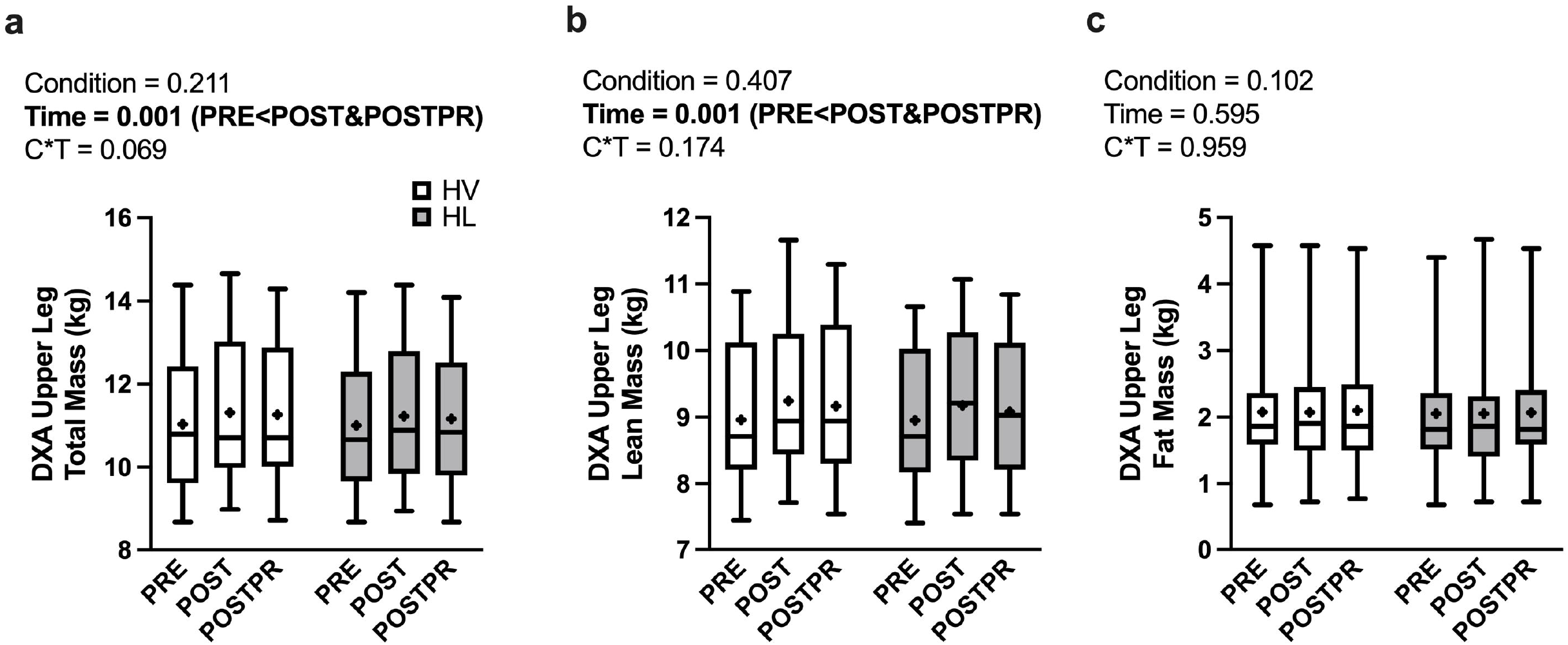
Segmental Upper Leg Composition. Data are presented as box and whiskers plots including median (central horizontal line), 25^th^ and 75^th^ percentile (box), minimum and maximum values (vertical lines) and mean values (cross) for DXA upper leg total mass (panel a), DXA upper lean mass (panel b), and DXA upper leg fat mass (panel c). Abbreviations: HV, high-volume; HL, high-load; DXA, dual energy x-ray absorptiometry.

### 2.5 Vastus Lateralis Muscle Morphology

A condition×time interaction was observed for magnetic resonance image (MRI)-derived VL cross-sectional area (*p*=0.046, η_p_^2^=0.211, Fig. 3a); however, no main effects of condition (*p*=0.490, η_p_^2^=0.037) or time (*p*=0.351, η_p_^2^=0.077) were observed. Post hoc analysis revealed no differences between conditions at PRE (*p*=0.246), POST (*p*=0.673), or POSTPR (*p*=0.247). There was no condition×time interaction (*p*=0.338, η_p_^2^=0.075, Fig. 3d) or main effect of condition (*p*=0.457, η_p_^2^=0.040) observed for ultrasound measured VL thickness. VL thickness changed over time (*p*=0.035, η_p_^2^ =0.241) where POST values were greater than PRE (*p*=0.026) and POSTPR (*p*=0.003). No differences were observed between PRE and POSTPR (*p*=0.614). There were no interactions observed for muscle pennation angle of the VL (*p*=0.393, η_p_^2^=0.064, Fig. 3f) or estimated VL muscle fiber length (*p*=0.602, η_p_^2^=0.036, Fig. 3c). Additionally, there were no main effects of condition or time for the aforementioned variables (*p*>0.05).

**Figure 3.**
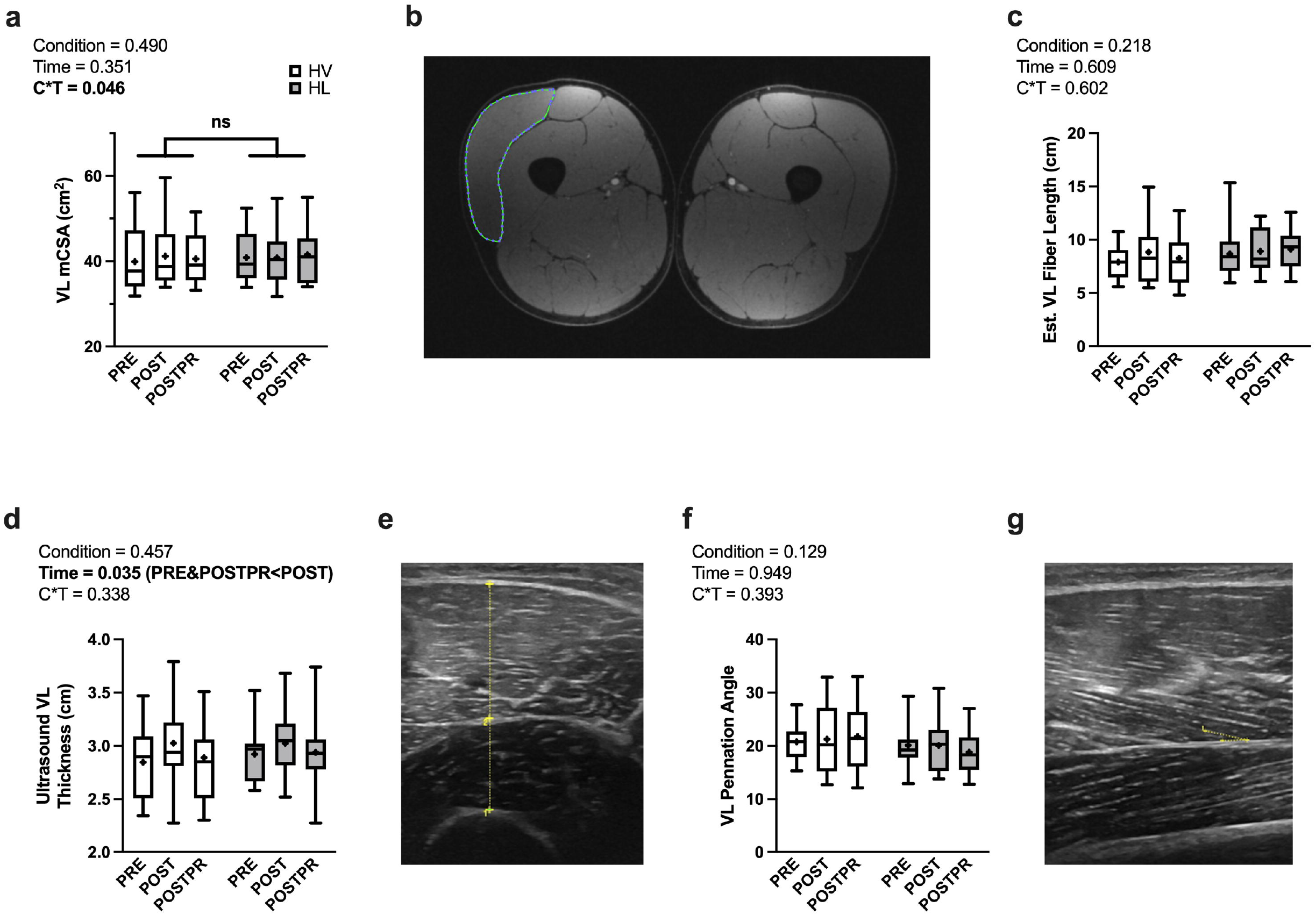
Vastus Lateralis Muscle Morphology. Data are presented as box and whiskers plots including median (central horizontal line), 25^th^ and 75^th^ percentile (box), minimum and maximum values (vertical lines) and mean values (cross) for VL mCSA (panel a), Est. VL fiber length (panel c), VL thickness (panel d), and VL muscle pennation angle (panel f). Representative images: Dual leg MRI for VL mCSA (panel b), ultrasound cross-section for VL thickness (panel e), ultrasound cross-section for pennation angle (panel g). No significance was observed following decomposition of condition×time interaction for VL mCSA. Abbreviations: HV, high-volume; HL, high-load; VL, vastus lateralis; mCSA, muscle cross-sectional area, Est., estimated.

### 2.6 Muscle Protein Adaptations

There was no condition×time interaction observed for sarcoplasmic protein concentrations per mg of wet tissue weight (*p*=0.112, η_p_^2^=0.159, Fig. 4a). There was a main effect of condition (*p*=0.002, η_p_^2^ =0.497, Fig. 4a) where sarcoplasmic protein concentrations in the HV group were higher than the HL group (44.8±1.6 versus 42.6±1.3 respectively). Additionally, there was a main effect of time (*p*= 0.022, η_p_^2^=0.239, Fig. 4a) where PRE sarcoplasmic protein concentrations were higher than POST (*p*=0.038) and POSTPR (*p*=0.032). No differences in sarcoplasmic protein concentrations were observed between POST and POSTPR (*p*=0.524)

**Figure 4.**
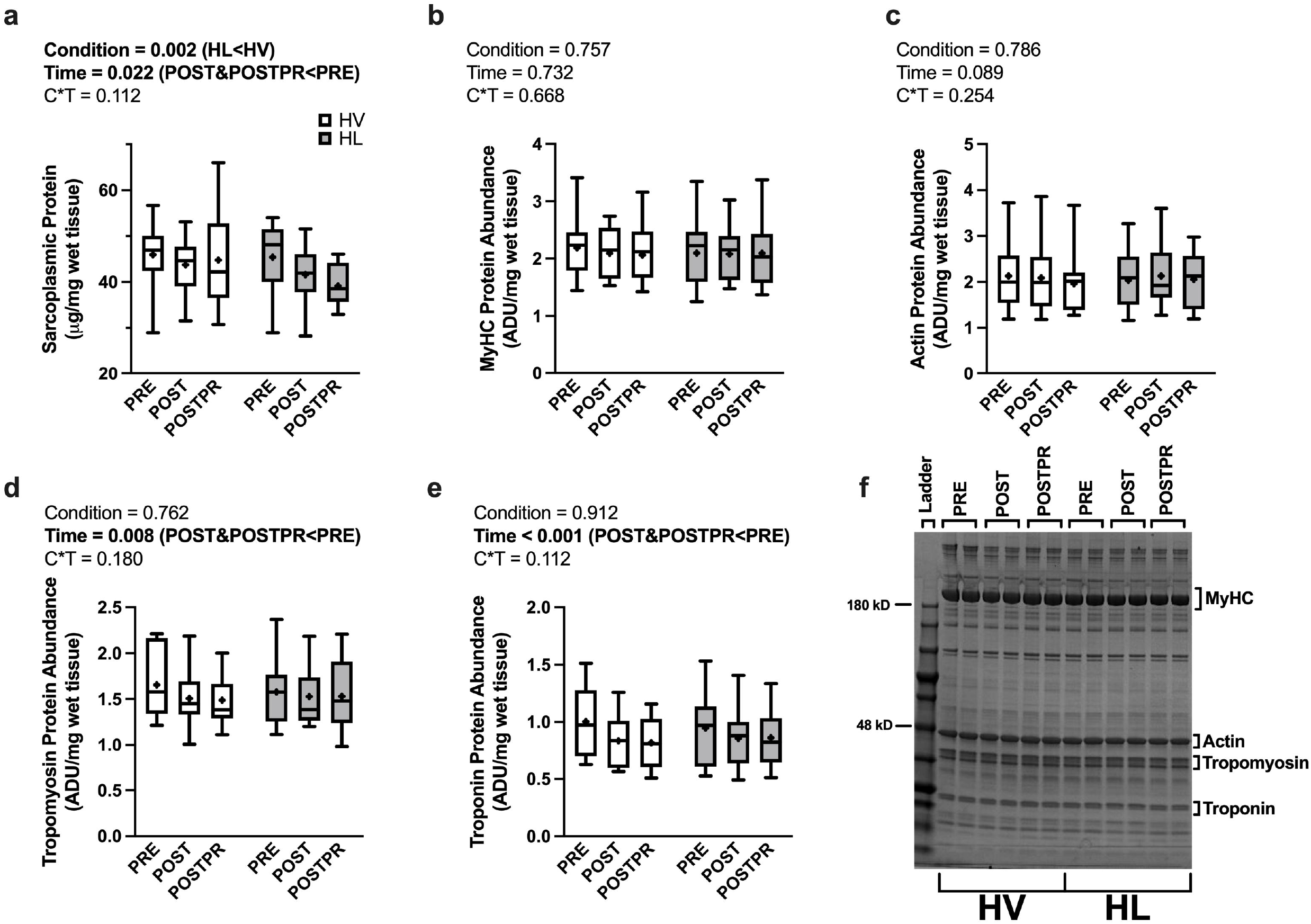
Muscle Protein Adaptations. Data are presented as box and whiskers plots including median (central horizontal line), 25^th^ and 75^th^ percentile (box), minimum and maximum values (vertical lines) and mean values (cross) for sarcoplasmic protein concentrations (panel a), MyHC protein abundance (panel b), actin protein abundance (panel c), tropomyosin protein abundance (panel d), and troponin protein abundance (panel e). Representative image: Coomassie blue stained poly-acrylamide gel for protein abundance. Abbreviations: HV, high-volume; HL, high-load; VL; MyHC, myosin heavy chain; ADU, arbitrary density units; kD, kilodalton.

There were no condition×time interactions observed for myosin heavy chain (MyHC) protein abundance per mg of wet tissue weight (*p*=0.668, η_p_^2^ =0.028, Fig. 4b) or actin protein abundance per mg of wet tissue weight (*p*=0.254, η_p_^2^=0.093, Fig. 4c). Additionally, no main effects of condition or time were observed for these variables (*p*>0.05). There was no condition×time interaction (*p*=0.180, η_p_^2^ =0.115, Fig. 4d) or main effect of condition (*p*=0.762, η_p_^2^=0.007, Fig. 4d) observed for tropomyosin protein abundance per mg wet tissue weight. However, a main effect of time was observed for this variable (*p*=0.008, η_p_^2^=0.294, Fig. 4d) where PRE was greater than POST (*p*=0.009) and POSTPR (*p*=0.010). No differences were observed between POST and POSTPR (*p*=0.704). There was no condition×time interaction (*p*=0.112, η_p_^2^=0.145, Fig. 4e) or a main effect of condition (*p*=0.912, η_p_^2^=0.001, Fig. 4e) observed for troponin protein abundance per mg wet tissue weight. A main effect of time was observed for this variable (*p*<0.001, η_p_^2^ =0.431, Fig. 4e) where PRE was greater than POST (*p*<0.001) and POSTPR (*p*=0.005). No differences were observed between POST and POSTPR (*p*=0.867).

### 2.7 Six-week Integrated Myofibrillar and Sarcoplasmic Protein Synthesis

No difference was observed in integrated myofibrillar protein synthesis (iMyoPS) between the HV and HL conditions (*p*=0.687; *d*=-0.106; Figure 5a). A significant difference was observed for integrated sarcoplasmic protein synthesis (iSarcoPS) where the HV condition exhibited a higher iSarcoPS (%/day) than the HL condition (*p*=0.0018; *d*=0.693; Figure 5b).

**Figure 5.**
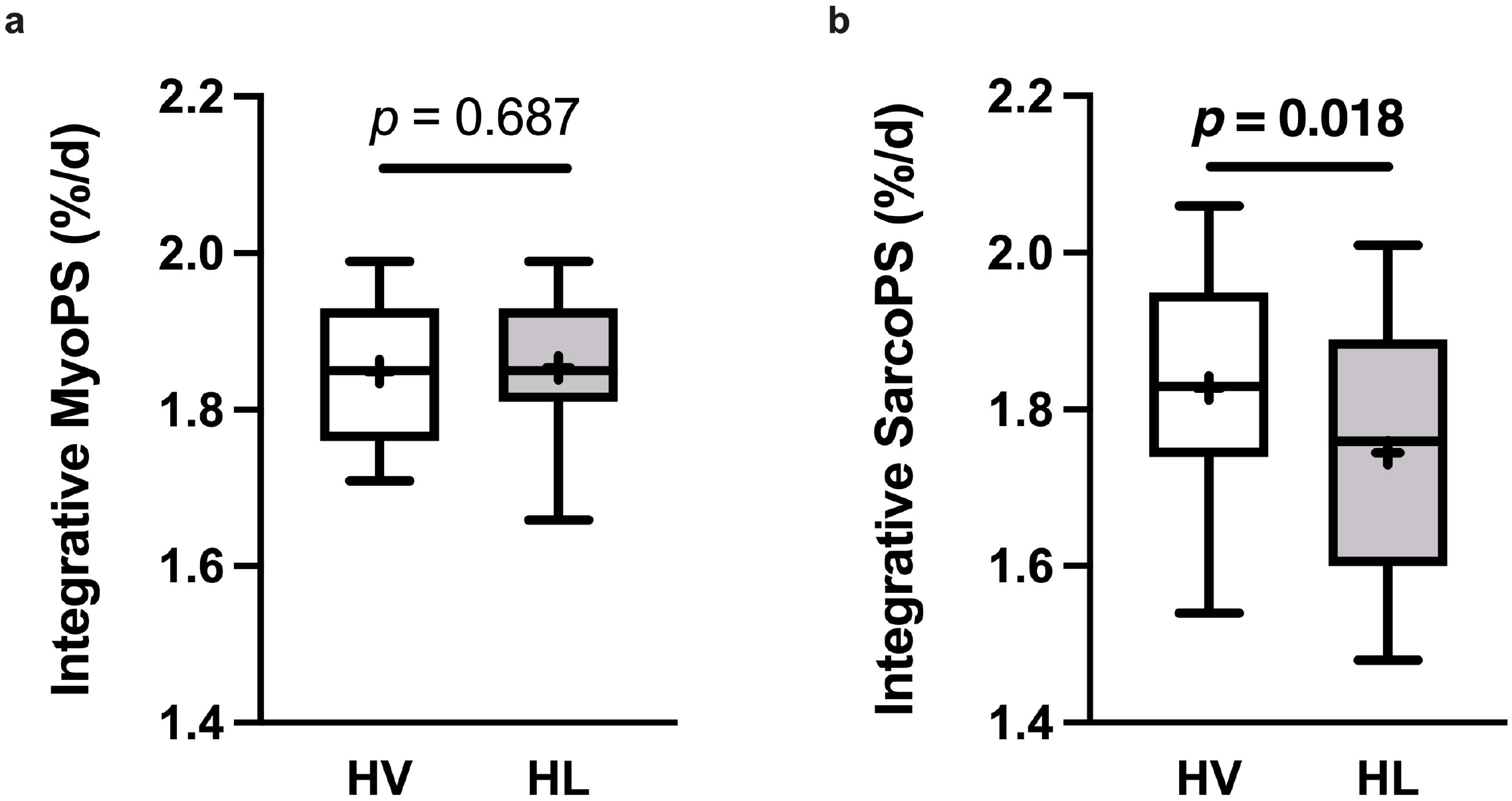
Six-week Integrated Myofibrillar and Sarcoplasmic Protein Synthetic Rates. Data are presented as box and whiskers plots including median (central horizontal line), 25^th^ and 75^th^ percentile (box), minimum and maximum values (vertical lines) and mean values (cross) for iMyoPS (panel a) and iSarcoPS (panel b). No significant differences were observed for iMyoPS between conditions. iSarcoPS was significantly lower in the HL condition as compared to the HV condition. Abbreviations: HV, high-volume; HL, high-load; MyoPS, myofibrillar protein synthesis; SarcoPS, sarcoplasmic protein synthesis.

## 3. DISCUSSION

Chief findings from the current study include: i) a condition×time interaction for MRI-derived VL mCSA although, when the model was statistically decomposed, no differences were found between conditions or over time, ii) an interaction for estimated unilateral leg extension 1RM where values were higher in the HL condition at POST and POSTPR when compared to the HV condition at the same time points, and iii) no significant difference in iMyoPS rates, albeit iSarcoPS was greater in the HV versus HL condition over the duration of the study. The relevance of these as well as other findings are discussed below.

The literature regarding macro-level muscle tissue changes following high- and low-load training are sparse. Furthermore, limitations in the available literature exist due to lack of congruency between loading paradigms and discrepancies on what constitutes low-load/high-volume and high-load/low-volume training. Nonetheless, there is prior literature that has interrogated differences between such training paradigms. Holm and colleagues^26^ reported that high-load (~70% 1RM) versus very low-load (~15.5% 1RM) leg extensor training increased quadriceps CSA; however, the change in the high-load condition was greater than the change in the low-load condition. Chestnut and Docherty^27^ reported similar increases in muscle CSA of the upper arm following 10 weeks of upper body resistance training using ~85% of 1RM for 6 sets of 4 repetitions versus ~70% for 3 sets of 10 repetitions. The aforementioned study by Mitchell and colleagues reported that performing three sets of knee-extensor training to fatigue at 30% or 80% of 1RM resulted in similar increases in quadriceps volume measured by MRI. Both modalities yielded greater quadriceps hypertrophy than performing one set at 80% 1RM to voluntary failure^12^. Furthermore, a systematic review and meta-analysis conducted by Schoenfeld and colleagues concluded that similar skeletal muscle growth can be realized across a variety of loading ranges^28^. As mentioned previously, while we observed a condition×time interaction for VL mCSA, decomposition of the model yielded no significant differences between HL and HV training. This is in agreement with previous literature showing that (on average), when loading exceeds 30% 1RM and training is executed near or at volitional fatigue, similar increases in whole-muscle hypertrophy can be realized.

Unilateral leg press and leg extension strength metrics increased over time, albeit there was no interaction observed for leg extensor peak torque or leg press strength changes. However, an interaction was observed for leg extension strength changes, where values increased more in the HL versus HV condition. Several studies have examined changes in strength between different loading paradigms. Campos and colleagues reported high-load resistance training (3-5RM) over an 8-week period yielded greater leg extension strength increases compared to high-volume resistance training (20-28RM)^29^; however, no differences in strength adaptations were reported between the 3-5RM group and a third group which performed training using 9-11RM loads. Additionally, Jenkins et al. published two studies comparing 30% 1RM versus 80% 1RM leg extensor training^14,15^. Results from both studies suggest that higher-load training elicited greater strength increases due to neural factors. Jessee and colleagues^30^ reported that unilateral training (4 sets to volitional failure) over an 8-week period resulted in greater strength adaptations for HL training (70% 1RM/no blood flow restriction) than low load conditions with or without blood flow restriction. Furthermore, Schoenfeld and colleagues reported increased barbell back squat strength with lower (30-50% 1RM) and higher-load (70-80% 1RM) training, with higher-load training resulting in greater strength adaptations^17^. When considering our findings in the context of these studies, it seems plausible that training at 60-90% 1RM over shorter-term periods may elicit similar strength changes. Collectively, our findings in the context of prior literature suggest that, while training at 30% 1RM seemingly yields similar increases in hypertrophy compared to training at higher loads, 30% 1RM training may not optimize strength gains due to differential neural adaptations.

A novel aspect of the current study was to compare how HL versus HV training affected molecular markers from muscle biopsies. This interrogation was prompted by select literature suggesting that a disproportionate increase in non-contractile proteins in myofibers may occur following high-volume resistance training. However, as reviewed by Jorgenson and colleagues, it should be noted that a large portion of the literature has shown that mechanical-load induced skeletal muscle hypertrophy is largely attributed to proportional increases in the contractile and non-contractile elements of the myofiber^31^. Nonetheless, a handful of studies exist showing that a disproportionate increase in non-contractile proteins may occur following months to years of resistance training^20,21,32,33^. Recently, our laboratory has reported increases in sarcoplasmic protein concentrations with concomitant decreases in the relative abundances of myosin heavy chain and actin protein abundances per mg of dry tissue weight following 6 weeks of extremely high-volume resistance training in previously-trained college-aged men^19^. We concluded at the time that “sarcoplasmic hypertrophy” may have occurred and that both of these molecular observations were reflective of this phenomenon. Our laboratory subsequently reported that sarcoplasmic protein concentrations were maintained while minor decrements occurred in actin protein abundance in previously-trained college-aged males that partook in a 10-week low-volume, high-load training paradigm^24^. When considering the findings from both studies, we hypothesized that HV training might facilitate sarcoplasmic hypertrophy, whereas HL training may facilitate proportional accretion of contractile and sarcoplasmic proteins with whole-muscle hypertrophy (i.e., conventional hypertrophy). In the current study, we observed decreased sarcoplasmic protein concentrations for both conditions at POST. Additionally, no significant changes in the relative abundances of actin and myosin heavy chain proteins were observed in either condition. Although these data disagree with prior findings from Haun et al., it is important to note the key differences that exist between the high-volume components of each study. First, Haun et al. utilized a bilateral exercise paradigm using the barbell back squat as the primary knee-extensor exercise, whereas the current study was a unilateral study design using a combination of unilateral leg press and leg extension to overload the knee-extensor muscles. To this end, it could be posited that training stress on the legs was much greater in the Haun et al. study versus the current study. Second, Haun et al. used a 6-week intervention starting at 10 sets of 10 repetitions per week (for each exercise) and finishing with 32 sets of 10 repetitions per week where loads were standardized at 60% 1RM^34^. The current study started the HV leg at 10 sets of 10 repetitions per week (split between two exercises) at week 1 and finished with 20 sets of 10 repetitions per week at week 6 where loads were standardized at 60% 1RM. Thus, although the HV leg was exposed to more training volume compared to the HL leg herein, the HV leg did not experience nearly the amount of volume as both legs incurred in the study by Haun et al. Moreover, the total training volume data in Figure 1 indicates that the HV leg was only exposed to ~11% more volume compared to the HL leg. We speculate that similar molecular adaptations between legs may have been due a relatively small difference in total training volume between legs throughout the duration of the study. Alternatively stated, a higher total training volume may have been needed to yield data suggestive of sarcoplasmic versus myofibrillar expansion in the HV condition. In spite of our null findings, however, it is intriguing that HV training increased iSarcoPS versus HL training. This partially supports the notion that HV training may affect the sarcoplasmic protein pool to a greater extent than HL training, but this warrants continued research.

As with many studies examining the effects of training interventions, the present study is limited due to a small sample size. The procurement of skeletal muscle tissue via percutaneous muscle biopsy inherently has a finite tissue yield. Due to this limitation, we lacked sufficient tissue to perform histology and assess fCSA values in each leg over time, and these assays would have been greatly beneficial for this study. While protein synthesis rates were measured herein, it is notable that muscle protein breakdown rates were not assessed and could have influenced outcomes. Additionally, previous literature has shown the ability to measure gross changes in skeletal muscle growth after 3-6 weeks of resistance training in untrained to recreationally trained men^34–36^. In this regard, we posit that the training status of the cohort in the current study may have precluded our ability to detect any meaningful training adaptations over the 6-week training period.

In conclusion, similar changes in muscle volume, morphology, and protein composition appeared with 6 weeks of HV versus HL training. However, HL training resulted in a greater increase in leg extension strength and higher 6-week iSarcoPS rates. Additionally, the current data challenge our prior muscle-molecular findings given: i) the lack of change in sarcoplasmic protein concentrations observed in the HV condition, and ii) no alterations being observed in myosin heavy chain and actin protein abundances following training. However, the current iSarcoPS findings suggest some muscle-molecular differences exist between HV and HL training, and warrant further research.

## 4. MATERIALS AND METHODS

### 4.1 Ethical approval and Pre-screening

Prior to study initiation, this protocol was reviewed and approved by the Auburn University Institutional Review Board and was conducted in accordance to the standards set by the latest revision of the Declaration of Helsinki (IRB approval #: 19-245 MR 1907).

College-aged resistance-trained men from the local community were solicited to participate in this study and were screened 4-7 days prior to the start of the study. Participants had to be free of cardio-metabolic diseases (e.g., morbid obesity, type II diabetes, severe hypertension), or any conditions that preclude the collection of a skeletal muscle biopsy. Additionally, training status for participants was determined by two criteria: i) self-reported resistance training >1 year at least 3 times weekly and; ii) a tested barbell back squat of ≥1.5× bodyweight (estimated from a 3 repetition maximum [3RM] test) in accordance to standards designated by the National Strength and Conditioning Association^37^ At the conclusion of the screening visit, participants were asked to maintain their current nutritional practices and to cease all training outside of the study.

### 4.2 Study design

A schematic of the study design is provided in Figure 6a. Briefly, participants performed a battery of testing prior to the start of training (PRE), 72 hours following the last bout of training after 6 weeks of unilateral lower body resistance training (POST), and 10 days following the last bout of training (POSTPR). The battery of tests performed are detailed further below, following a description of the training intervention and tracer methodologies.

**Figure 6.**
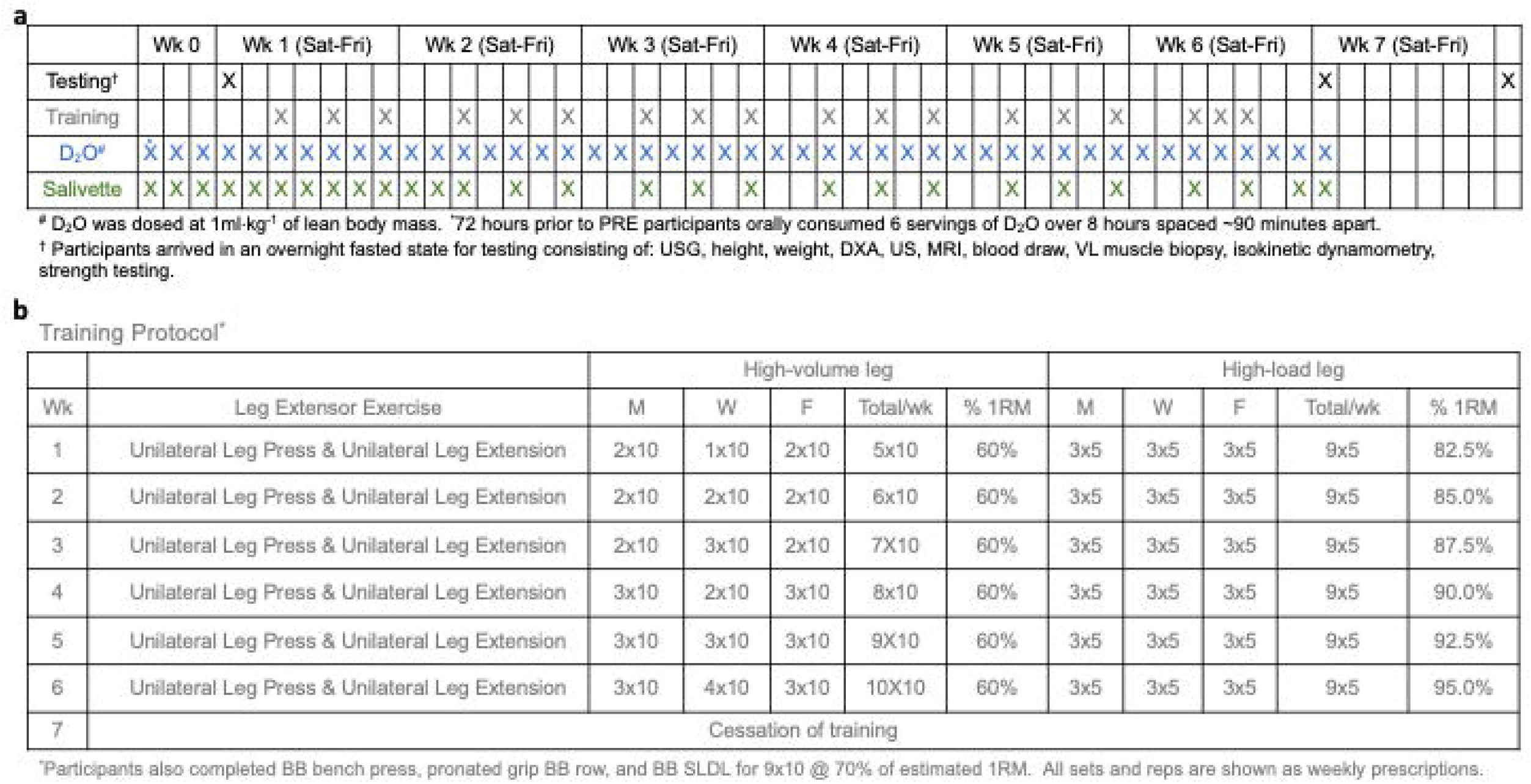
Study Design. Panel a provides an overview of testing, training, D2O administration, and saliva collection times. Panel b provides a schematic of training by day and total training for each week. Abbreviations: wk, week; D2O, deuterium oxide.

### 4.3 Resistance Training

Participants performed overloading unilateral lower body resistance training (i.e., singleleg leg press and single-leg leg extension) 3 days per week in conjunction with compound upper body exercises (i.e., barbell bench press, pronated grip barbell row, barbell stiff-leg deadlift). Notably, participants were randomly assigned to lower body training conditions prior to the start of the study, where some participants performed HV training on the left leg and HL training on the right leg or vice versa. All upper body exercises were performed for 3 sets of 10 repetitions at 70% of tested 1RM. Progression for the lower body training can be found in Figure 6b.

### 4.4 Isotope Tracer Protocol

Deuterium oxide (D_2_O) (Cambridge Isotope Laboratories, Inc.; Andover, MA, USA) was provided to the participants three days prior to and over the first 6 weeks of the study at 1 mL•kg^-1^ of lean body mass. For rapid enrichment of deuterium (^2^H) participants were instructed to orally consume 6 doses of D_2_O over an eight-hour period, three days prior to the first data collection (PRE), and were instructed to consume a top-up dose daily thereafter consisting of one dose of D_2_O until data collection was performed at the conclusion of week 6 of the study (POST). Saliva samples were taken utilizing sterile salivettes (SARSTEDT AG & Co, Nümbrect, Germany). Briefly, participants were instructed to chew on the cotton swab for 1 min and place the swab back into the top compartment of the salivette. This process was completed daily for the first 10 days of the study and every Monday, Wednesday, and Friday thereafter. Participants were instructed to place salivettes in their home freezers on days when saliva was donated outside of the laboratory. Samples were stored at −20 □ until further processing as described below.

### 4.5 Testing sessions

#### Urine specific gravity testing for adequate hydration

Upon arrival to each testing session participants submitted a urine sample (~5mL) for urine specific gravity (USG) assessment. Measurements were performed using a handheld refractometer (ATAGO; Bellevue, WA, USA). USG levels in all participants were ≤ 1.020 indicative of a euhydrated state^38^ and thus were considered adequately hydrated for further testing.

#### Body composition testing

Following hydration testing, participants underwent height and weight testing utilizing a digital column scale (Seca 769; Hanover, MD, USA) with body mass collected to the nearest 0.1 kg and height to the nearest 0.5 cm. Participants were then subjected to a full body dual energy x-ray absorptiometry (DXA) scan (Lunar Prodigy; GE Corporation, Fairfield CT, USA). Our laboratory^39^ has previously shown same day reliability of the DXA during test-calibrate-retest on 10 participants to yield an intra-class correlation coefficient (ICC) of 0.998 for total body lean mass.

#### Measurements of muscle morphology

Following body composition testing, participants were tested for VL muscle thickness and muscle pennation angle via ultrasound. VL thickness of both legs were assessed by placing a 3 to 12 MHz multi-frequency linear phase array transducer (Logiq S7 R2 Expert; General Electric, Fairfield, CT, USA) midway between the iliac crest and lateral epicondyle of the femur. Measurements were taken from a standing position and participants were instructed to bear the majority of their weight on the leg contralateral to the leg being measured. VL pennation angles were taken immediately following thickness measures by placing the transducer longitudinally at the same site mentioned above. VL thickness was measured as the distance between the superficial and deep aponeurosis while VL pennation angle was measured as the angle of the deep aponeurosis as it relates to the individual fascicles.

Estimated fiber length was calculated using methods similar to those described by Fukunaga and colleagues^40^.

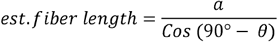

Whereas a is equal to the distance between the superficial fascia and the deep aponeurosis and *θ* is equal to the angle of pennation. Importantly – to minimize variability in measurements as suggested in previous studies^41,42^ – all measures were taken by the same investigator (S.C.O.) whom in a test-retest validation on 11 participants has been found to have an ICC of 0.983. Critically, this investigator was not privy to the (HV or HL) condition for each image. Moreover, the location of measurements were marked, using anatomical landmarks, by the investigator so that the subsequent MRI scans and muscle biopsies could be obtained from the same plane of measurement.

#### MRI for muscle cross-sectional area

Following ultrasound assessments, participants were shuttled to the Auburn University MRI Research Center to perform dual-leg mid-thigh MRI scans. All measurements were performed on a 3T VARIO system (Siemens, Erlangen, Germany). Briefly, upon arrival participants were placed in a supine position for 10 minutes to allow for body fluid stabilization to occur. Volume coil was used for RF transmit and body and spine array coils placed around the legs were used for signal receive. 3D gradient echo sequence (3D fast low angle shot) was used to acquire fat suppressed images with the following parameters: TR/TE = 10/4.92 ms; flip angle = 10°; bandwidth = 510 Hz/pixel, in-plane resolution 1mm X 1mm and slice thickness = 2.2mm. An axial 3D 35.2mm thick slab (a6 partitions) was placed to image both thighs with the thickness dimension carefully centered on the participant biopsy marking. Following the conclusion of the study, MRI scans were digitized offline using Osirix MD software (Pixmeo, Geneva, CHE), and software tools were used to manually trace the border of the VL yielding mCSA values. All MRI scans and image analyses were performed by the same investigators (R.J.B. and M.A.S., respectively), without knowledge of the (HV or HL) condition.

#### Collection of muscle tissue

Following MRI scans, right and left leg VL muscle biopsies were collected using a 5-gauge needle under local anesthesia as previously described^43,44^. Immediately following tissue procurement, tissue was teased of blood and connective tissue, wrapped in pre-labelled foils, flash frozen in liquid nitrogen, and subsequently stored at −80°C for molecular analyses described below.

#### Strength Testing

Following muscle skeletal muscle biopsies, participants underwent isokinetic dynamometry (Biodex System 4; Biodex Medical Systems, Inc., Shirly, NY, USA) for leg extensor peak torque and 3RM testing. For right and left leg extensor peak torque testing, participants were fastened to the isokinetic dynamometer. Each participant’s lateral epicondyle was aligned with the axis of the dynamometer, and seat height was adjusted to ensure the hip angle was approximately 90°. Prior to torque assessment, each participant performed a warm-up consisting of submaximal to maximal isokinetic knee extensions. Participants then completed five maximal voluntary isokinetic knee extension actions at 1.05 rad/s (60°/s) and 2.09 rad/s (120°/s). Participants were provided verbal encouragement during each contraction. The isokinetic contraction resulting in the greatest value was used for analyses. Following isokinetic dynamometry participants performed maximum strength testing for the exercises utilized over the duration of the study (single-leg leg press, single-leg leg extension, barbell bench press, pronated grip barbell row, and barbell stiff-leg deadlift). Briefly, participants performed 3 warmup sets starting at ~50% of their self-selected opening weight for 10 repetitions, then 75% of their self-selected opening weight for 5 repetitions, and 90% of their self-selected opening weight for 3 repetitions. Following warm-ups, participants executed their opening attempt for 3 repetitions with 5-10% increases being made from there on until a 3RM was achieved. All strength testing was performed by investigators holding the NSCA certified strength and conditioning specialist credential (C.G.V. and C.L.S.). This process was completed for all exercises at PRE while only the single-leg exercises were tested at POST and POSTPR.

### 4.6 Biochemical assays

#### Sarcoplasmic and myofibrillar protein isolation

Isolation of protein fractions was performed using the proteomic validated “MIST” or “myofibrillar isolation and solubilization technique”^45^. 1.7 mL polypropylene tubes were pre-filled with ice-cold buffer (300 μL; Buffer 1: 25 mM Tris, pH 7.2, 0.5% Triton X□100, protease inhibitors) and placed on ice. Skeletal muscle foils were removed from −80°C, placed on a liquid nitrogen-cooled ceramic mortar and pestle, and tissue was pulverized into 2-4 mm^3^ chunks. Chunks (~20 mg) were weighed using a scale with a sensitivity of 0.0001 g (Mettler-Toledo; Columbus, OH, USA) and placed into 1.7 mL polypropylene tubes with buffer and placed on ice. Samples were homogenized using tight-fitting pestles and centrifuged at 1,500 g for 10 minutes at 4°C. Supernatants (sarcoplasmic fraction) were collected and placed in new 1.7 mL polypropylene tubes on ice. As a wash step, the resultant myofibrillar pellet was resuspended in 300 μL of Buffer 1 and centrifuged at 1,500 g for 10 minutes at 4°C. The supernatant was discarded and the myofibrillar pellet was solubilized in 300 μL of ice-cold resuspension buffer (20 mM Tris-HCl, pH 7.2, 100 mM KCl, 20% glycerol, 1 mM DTT, 50 mM spermidine, protease inhibitors). Protein concentrations for the sarcoplasmic fraction were determined the same day as protein isolations to minimize freeze-thaw artifact, and the myofibrillar fraction was prepared for actin and myosin heavy chain protein abundance analyses and stored at −80°C until analysis occurred. The methodologies used for the above are described in further detail below.

#### Determination of sarcoplasmic protein concentrations

Sarcoplasmic protein resuspensions were batch-assayed for determination of protein concentration using a commercially available bicinchoninic acid (BCA) kit (Thermo Fisher Scientific; Waltham, MA, USA). Samples were assayed in duplicate (sarcoplasmic protein) using a microplate assay protocol where a small volume of sample was assayed (20 μL of 5x diluted sample + 200 μL Reagent A + B). The average duplicate coefficient of variation for sarcoplasmic protein concentration was 2.27%.

#### SDS-PAGE and Coomassie staining for relative contractile protein abundance

Determination of contractile protein abundances per mg wet tissue were performed as previously described by our laboratory and others^19,44,46,47^. Briefly, SDS-PAGE sample preps were made using 10 μL resuspended myofibrils, 65 μL distilled water (diH2O), and 25 μL 4x Laemmli buffer. Samples (5 μL) were then loaded on precast gradients (4-15%) SDS-polyacrylamide gels in duplicate (Bio-Rad Laboratories) and subjected to electrophoresis at 180 V for 40 minutes using pre-made 1x SDS-PAGE running buffer (Ameresco). Following electrophoresis, gels were rinsed in diH2O for 15 minutes and immersed in Coomassie stain (Lab Safe GEL Blue; G-Biosciences; St. Louis, MO, USA) for 2 hours. Gels were then destained in diH2O for 60 minutes, and band densitometry was performed with a gel documentation system and associated software (ChemiDoc; Bio-Rad Laboratories, Hercules, CA, USA). Given that a standardized volume from all samples was loaded onto gels, myosin heavy chain and actin band densities were normalized to input muscle weights to derive arbitrary density units (ADU) per mg wet muscle. All values were then divided by the mean of the PRE time point to depict myosin heavy chain, actin, tropomyosin, and troponin abundances. Our laboratory has reported that this method yields exceptional sensitivity in detecting 5-25% increases in actin and myosin content^44^. Average duplicate coefficients of variation for relative actin, myosin, tropomyosin, and troponin protein concentrations herein were 1.95%, 1.90%, 2.22%, and 3.54% respectively.

#### Six-week integrated myofibrillar and sarcoplasmic protein synthesis rates

Protein isolations were performed using ~30 mg of tissue utilizing the MIST method as described above. Prior to preparation for tracer analysis the sarcoplasmic protein fraction was lyophilized and precipitated in 1 mL of 1M perchloric acid to form a pellet. The myofibrillar pellet was purified by adding 500 μL of DDH_2_O followed by vortexing for 5 s and centrifugation at 1500 rpm at 4□ for 10 minutes. Following centrifugation, 1 mL of 0.3 M NaOH was added to the sample and then vortexed for 5 s followed by being placed in a heat block at 50□ for 30 minutes of which samples were vortexed for 5 s every 10 minutes. Samples then underwent centrifugation at 10,000 rpm at 4□ for 10 minutes. The supernatant (sarcoplasmic or myofibrillar fraction) was transferred into a 4 mL glass screw-top tube. 1 M perchloric acid was then added to tubes and centrifuged at 2500 rpm at 4□ for 10 minutes. The supernatant was removed, and the remaining pellet was washed in 70% ethanol and centrifuged at 2500 rpm at 4□ for 10 minutes twice. Amino acids were extricated through the addition of 1 mL of 1 Dowex resin (50WX8-200 resin; Sigma Aldrich) and 1 mL of 1 M HCl prior to heating at 110 ≥ for 72 hours. Cation exchange columns were used to isolate the free amino acids after which the amino acids were analyzed for deuterated-alanine content (^2^H-alanine) using a gas chromatography pyrolysis isotope ratio mass spectrometer. The amino acids were derivatized as their *n*□ methoxycarbonyl methyl esters. Dried samples were suspended in 60 μl distilled water and 32 μl methanol, and following vortex, 10 μl of pyridine and 8 μl of methylchloroformate were added. Samples were vortexed for 30 s and left to react at room temperature for 5 min. The newly formed *n*□ methoxycarbonyl methyl esters of amino acids were then extracted into 100 μl of chloroform. A molecular sieve was added to each sample for ~20 s before being transferred to a clean glass Gas Chromatography insert. Incorporation of deuterium into protein bound alanine was determined by gas chromatography-pyrolysis-isotope ratio mass spectrometry (Delta V Advantage) alongside a standard curve of known l alanine 2,3,3,3 d4 enrichment to validate measurement accuracy of the instrument^48^.

The water phase of saliva was injected 6 times with the average of the last 3 injections being used for data analysis. The H isotope enrichments for both muscle and saliva were initially expressed as δ^2^H% and then converted to atom percent excess using standard equations as reported by Wilkinson et al^48^. Fractional synthetic rate for the myofibrillar and sarcoplasmic protein fractions were calculated using the standard precursor-product method as described by other laboratories^49–51^.

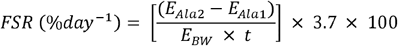

Whereas E_Ala1_ and E_Ala2_ represent ^2^H enrichment at PRE and POST respectively (in atom percent excess) from skeletal muscle biopsies. E_BW_ is the average ^2^H enrichment (in atom percent excess) of total body water between time-points and *t* is time in the number of days D_2_O was ingested. Multiplying by 3.7 adjusts for average ^2^H atoms that can be bound to alanine and multiplying by 100 converts this to a percentage per day^48,52^.

### 4.7 Statistical Analyses

Statistical analyses were performed using SPSS (Version 26; IBM SPSS Statistics Software, Chicago, IL, USA), open-source software JASP (Version 0.11.0; JASP Team; 2019), and RStudio (Version 1.1.463, R Foundation for Statistical Computing, Vienna, AT). Prior to analysis, assumptions testing for normality was performed using Shapiro-Wilk’s test for all dependent variables. If the assumption of heteroscedasticity was violated for repeated measures, a Greenhouse-Geisser correction factor was applied. Dependent variables were analyzed using multi-factorial repeated measures ANOVAs, and LSD *post hoc* tests were used to assess differences in dependent variables for leg or time. Statistical significance for null hypothesis testing was set at *p*<0.05. Data are presented throughout as mean ± standard deviation (bar graphs) or box and whiskers plots including median (central horizontal line), 25^th^ and 75^th^ percentile (box), minimum and maximum values (vertical lines) and mean values (cross).

## AUTHOR CONTRIBUTIONS

CGV and MDR designed the study and primarily drafted the manuscript. PWM and CGV ran statistical analyses. CGV designed the training program and coordinated the study. All other authors assisted with testing, assays, or other aspects of the studies.

## ACKNOWLEDGEMENTS

The authors would like to thank the participants for their dedication to executing this study. We would also like to thank Johnathon Moore, Samantha Slaughter, Andy Cao, Max Coleman, Max Michel, Megan Edwards, and Sullivan Clement for their assistance in collecting data.

